# Cytos MPS: an automated multimodal pooled screening platform for protein, morphology and transcriptomics

**DOI:** 10.64898/2026.09.20.752925

**Authors:** Tyler Lopez, John Rabalais, Ben Song, Andrew Boddicker, Ashley Schwartz, Daniel Honigfort, Xiaodong Qi, Junhua Zhao, Matthew Kellinger, Semyon Kruglyak

**Affiliations:** Element Biosciences, San Diego, CA, USA

## Abstract

Cytos Multimodal Pooled Screening (Cytos MPS) measures morphology, protein abundance and localization, CRISPR guide identity, and native 3′ transcriptome in the same fixed cells. Following IL-1β stimulation, cells with higher nuclear NF-κB show lower phospho-p38/HSP27 and higher NFKBIA/DUSP1 in the same cell, a measurement neither imaging nor transcriptomics alone provides. TGFBR2 and CTNNB1 perturbations produce distinct multimodal phenotypes. Guide representation and perturbation transcriptional responses agree with an orthogonal single-cell transcriptomic platform.

## Main

Linking a genetic perturbation to its cellular consequences increasingly requires more than one molecular readout. Optical pooled screening (OPS) links guide identity to imaging phenotypes, including protein localization^1^, and Cell Painting extends morphological profiling^2^. Perturb-seq profiles transcriptional responses in dissociated, pooled-perturbation cells^3^. Spatial approaches retain cellular context: Perturb-FISH combines guide identity with spatial transcriptomics, without a matched protein panel^4^; spatial CRISPR genomics (Perturb-map) maps guide identity onto tissue histology and transcriptomes, also without a matched protein panel^5^. Cytos MPS combines all four: morphology, immunofluorescence protein abundance and localization, CRISPR guide identity, and native 3′ transcriptome, from the same fixed cell, using Direct In-Sample Sequencing^6^ to read CRISPR guides and 3′ transcripts without PCR. All four modalities are registered to a common per-cell key in one spatial data object.

We applied the platform to A549 cells carrying a pooled six-gene CRISPR knockout library, in two independent runs (241,730; 239,725 cells; 12-well slides, untreated and IL-1β-stimulated at 15 and 30 minutes). Paired values throughout are given as run 1; run 2. The platform assigned 85.9%; 87.0% of cells to a single guide, yielded PCR-free molecule counts averaging 1,626; 1,630 per cell, and detected 12,163; 12,237 genes above a false-positive-calibrated threshold (Fig. 1).

**Figure 1.**
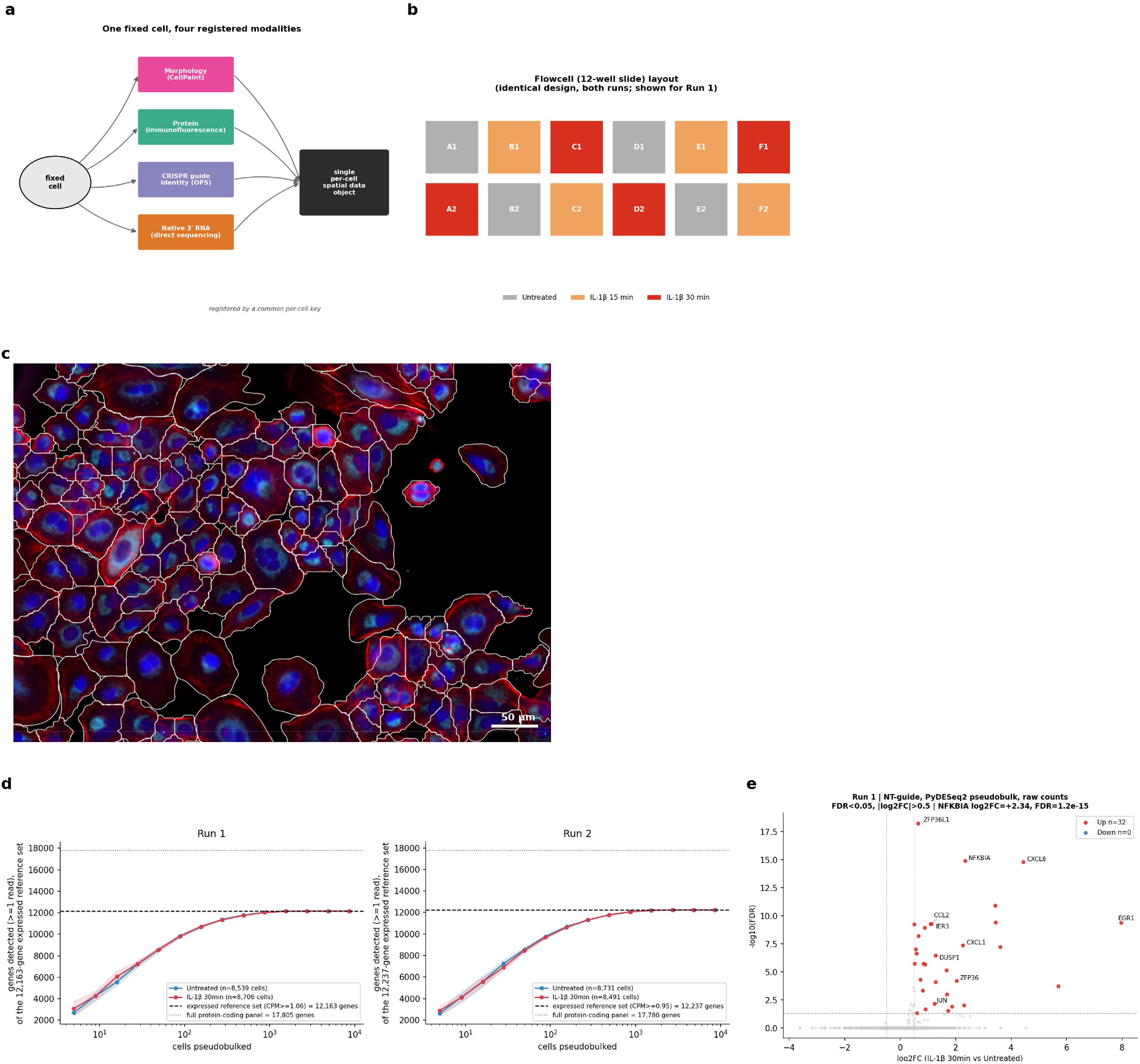
Platform overview and capability metrics. **a**, Morphology, protein, guide identity and native 3′ RNA from the same fixed cell, in acquisition order, registered by a common per-cell key into one spatial data object. **b**, 12-well slide layout, identical in both runs: untreated, IL-1β 15 min and IL-1β 30 min, four wells each. **c**, CellPaint image of an untreated well, run 1: nuclei (blue), actin (red) and mitochondria (cyan), with per-cell segmentation outlines (white). Per-run QC (run 1; run 2): 241,730; 239,725 cells; 85.9%; 87.0% assigned to a single guide; 99.6%; 99.6% nucleated; 78.9%; 79.3% confluency. **d**, Gene-detection saturation curves for untreated (blue) and IL-1β 30-min (red) non-targeting cells, both runs; mean ± s.d.; 10x comparison in Extended Data Fig. 10. **e**, IL-1β 30-min vs. untreated differential expression in non-targeting cells, run 1 (PyDESeq2 pseudobulk; FDR < 0.05, |log2FC| > 0.5); 32 induced genes, none repressed.

We chose IL-1β stimulation as the test system because its pathway is fast and well characterized, its activation is an imageable event (NF-κB p65 nuclear translocation) with known transcriptional targets, and its receptor, IL1R1, was among the knockouts in the library. We first asked whether the platform readouts reproduce this known IL-1β/NF-κB biology^7^. Non-targeting cells showed the expected nuclear translocation of NF-κB p65 upon stimulation (Cohen’s d = +2.76; +2.71, both P < 10^−5^; Fig. 2a,b), and IL1R1 knockout attenuated the mean translocation response by 90% in both runs (P = 2.1×10^−5^; 9.5×10^−7^), with matching transcriptional blunting: 32; 39 stimulation-induced genes in non-targeting cells, led by NFKBIA (log2FC +2.34) and CXCL8, with all 12 top induced genes showing a smaller fold-change under IL1R1 knockout (Fig. 1e; Extended Data Figs. 1, 2). A subset of IL1R1-knockout cells nonetheless retained elevated nuclear NF-κB, disproportionately attributable to a single guide (IL1R1_4; Extended Data Fig. 3). Matched protein and RNA measurements thus captured attenuation of both signaling and transcriptional responses.

**Figure 2.**
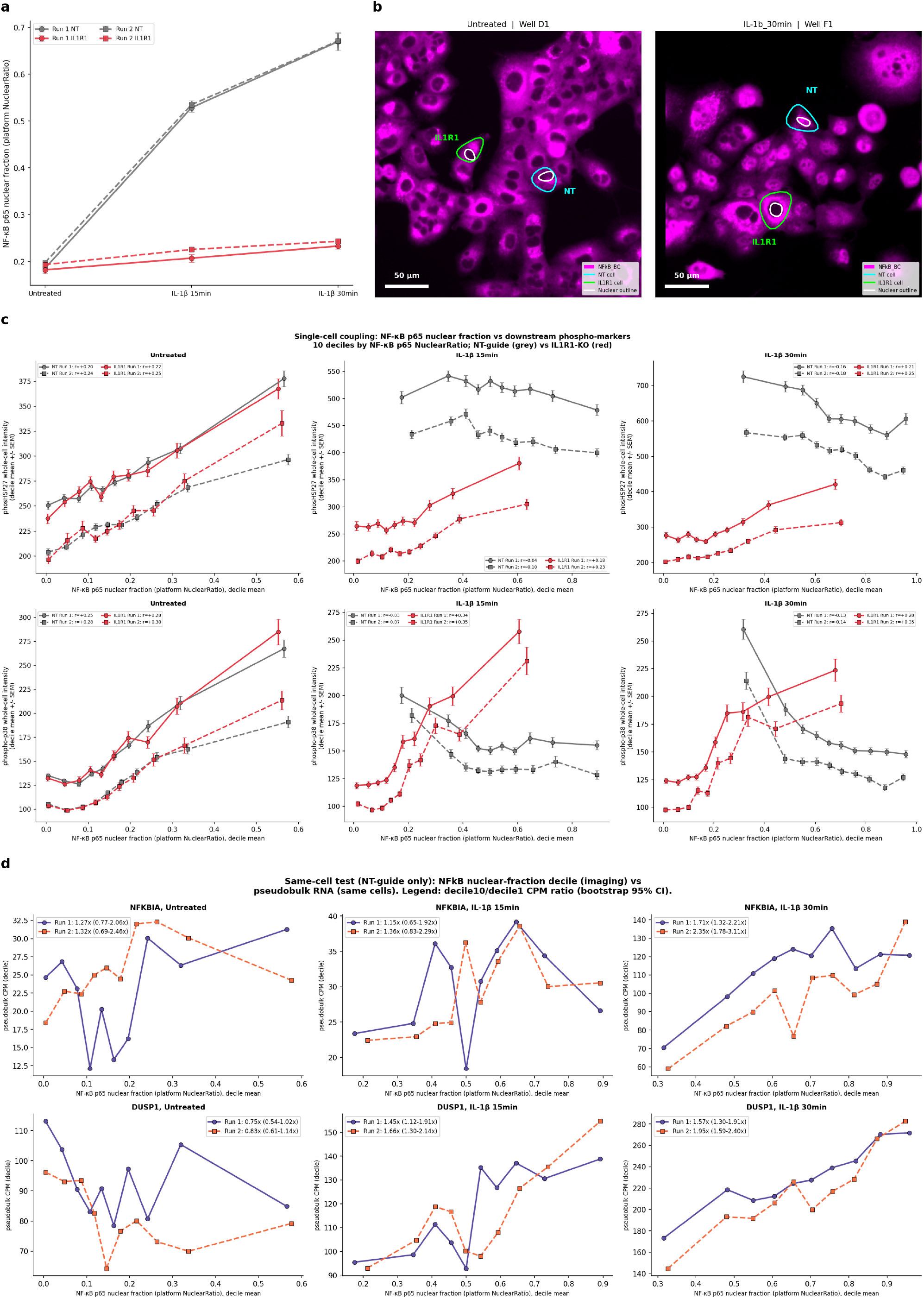
IL1R1 knockout reproduces known IL-1β/NF-κB biology; same-cell resolution reveals receptor-dependent NF-κB/MAPK/RNA coupling. **a**, NF-κB nuclear fraction timecourse, per-well means ± s.e.m.; non-targeting (grey) vs. IL1R1-KO (red); marker distinguishes run. **b**, Representative cells (NF-κB p65 channel), untreated vs. IL-1β 30-min; cyan/green outlines, non-targeting/IL1R1-KO cells; white, nuclear boundary; scale bars, 50 µm. **c**, Decile-mean phospho-p38 and phospho-HSP27 intensity vs. decile-mean NF-κB nuclear fraction, non-targeting vs. IL1R1-KO, both runs; Spearman’s ρ on per-cell values annotated. **d**, Pseudobulk NFKBIA and DUSP1 across the same deciles, non-targeting cells; decile-10:decile-1 CPM ratio (95% cell-bootstrap CI) in legend.

We next asked whether a cell’s NF-κB activation predicts its own downstream signaling and transcription. Because translocation is scored per cell, the same cells’ MAPK phospho-signaling and RNA state can be examined at single-cell resolution, a joint measurement neither imaging alone (no transcriptome) nor a dissociated transcriptomic screen (ref. 3; no matched morphology/protein channel) can make. In stimulated non-targeting cells, phospho-p38 and phospho-HSP27 were negatively associated with NF-κB nuclear fraction at 30 minutes (ρ = −0.13; −0.14 and −0.16; −0.18, respectively). This relationship strengthened after cell-area adjustment (area-adjusted ρ = −0.22; −0.21 for phospho-p38), whereas IL1R1 knockout and untreated non-targeting cells showed positive associations (Fig. 2c). Across NF-κB nuclear-fraction deciles, NFKBIA and DUSP1 expression increased at 30 minutes (decile-10:decile-1 CPM ratios 1.71; 2.35 and 1.57; 1.95, respectively). Both phospho-marker correlations were negative and both RNA ratios exceeded 1 in all four wells of each run at 30 minutes, including with RNA deciles defined within wells (Methods). DUSP1 increased at 15 minutes (ratio 1.45; 1.66), whereas NFKBIA’s cell-bootstrap confidence interval still spanned 1. Untreated cells showed the opposite DUSP1 trend (ratio 0.75; 0.83; Fig. 2d); CXCL8/ZFP36L1 relationships were weaker (Extended Data Fig. 4).

Depth controls and comparison with ∼1,600–1,700 background genes argue against generic technical or cell-activity effects explaining the RNA associations: NFKBIA/DUSP1 exceeded 91.1–99.9% of the background ratios (Extended Data Fig. 5). The timing is consistent with delayed negative feedback through DUSP1-mediated p38 dephosphorylation^8^ (Extended Data Fig. 6), although differing pathway kinetics remain an alternative explanation. Matched protein and RNA measurements thus suggest how transcriptional induction could contribute to attenuation of phospho-signaling.

A separate pair of constitutive knockouts, TGFBR2 (TGF-β receptor) and CTNNB1 (Wnt/β-catenin effector), allowed us to characterize perturbation-associated changes across modalities at baseline. TGFBR2 and CTNNB1 knockout produced opposite cell-geometry phenotypes across all four measured features (cell area, major and minor axis, nuclear-area fraction): TGFBR2 cells were smaller with a higher nuclear-area fraction (cell area −11.8%; −10.9%, paired t-test, n = 4 wells/run), while CTNNB1 cells were larger with a lower nuclear-area fraction (Fig. 3a). Effects were modest (|d| = 0.08–0.21) but reversed in sign between genotypes in both runs. Each knockout’s geometry change was accompanied by broader CellPaint intensity changes (36; 31 of 40 features for TGFBR2, 38; 37 for CTNNB1, FDR < 0.05), led in TGFBR2 by cell-membrane signal (d = +0.45; +0.47; Extended Data Fig. 7a). Baseline differential expression also distinguished the genotypes (TGFBR2: 113; 71 genes, including reduced membrane/adhesion transcripts ITGAV, CDH2, CLDN2, ANTXR1, concordant both runs; CTNNB1: 156; 171 genes; Fig. 3b; Extended Data Figs. 8, 9), and the protein panel shifted in both knockouts (TGFBR2: 125; 133 of 160 features, led by NF-κB p65 and Na^+^/K^+^-ATPase up, alpha-tubulin down; CTNNB1: 133; 130 of 160 features; Extended Data Fig. 7b for full per-feature profiles). The NF-κB p65 protein shift itself persisted after adjustment for cell area (Cohen’s d changed by less than 20% after regressing out area); TGF-β and Wnt signaling have each been reported to cross-talk with NF-κB, though direction is context-dependent^9^,^10^. The remaining knockouts served as specificity controls at baseline: OR1I1 (0 significant genes, both runs) and GSK3B (1, itself only, both runs) showed essentially no transcriptional effect, arguing against a generic knockout artifact, while SMAD7 induced a small, reproducible signature (1; 3 significant genes) driven in both runs by the same gene, ID1, a canonical BMP/TGF-β-superfamily target de-repressed by loss of its pathway inhibitor (Extended Data Fig. 9).

**Figure 3.**
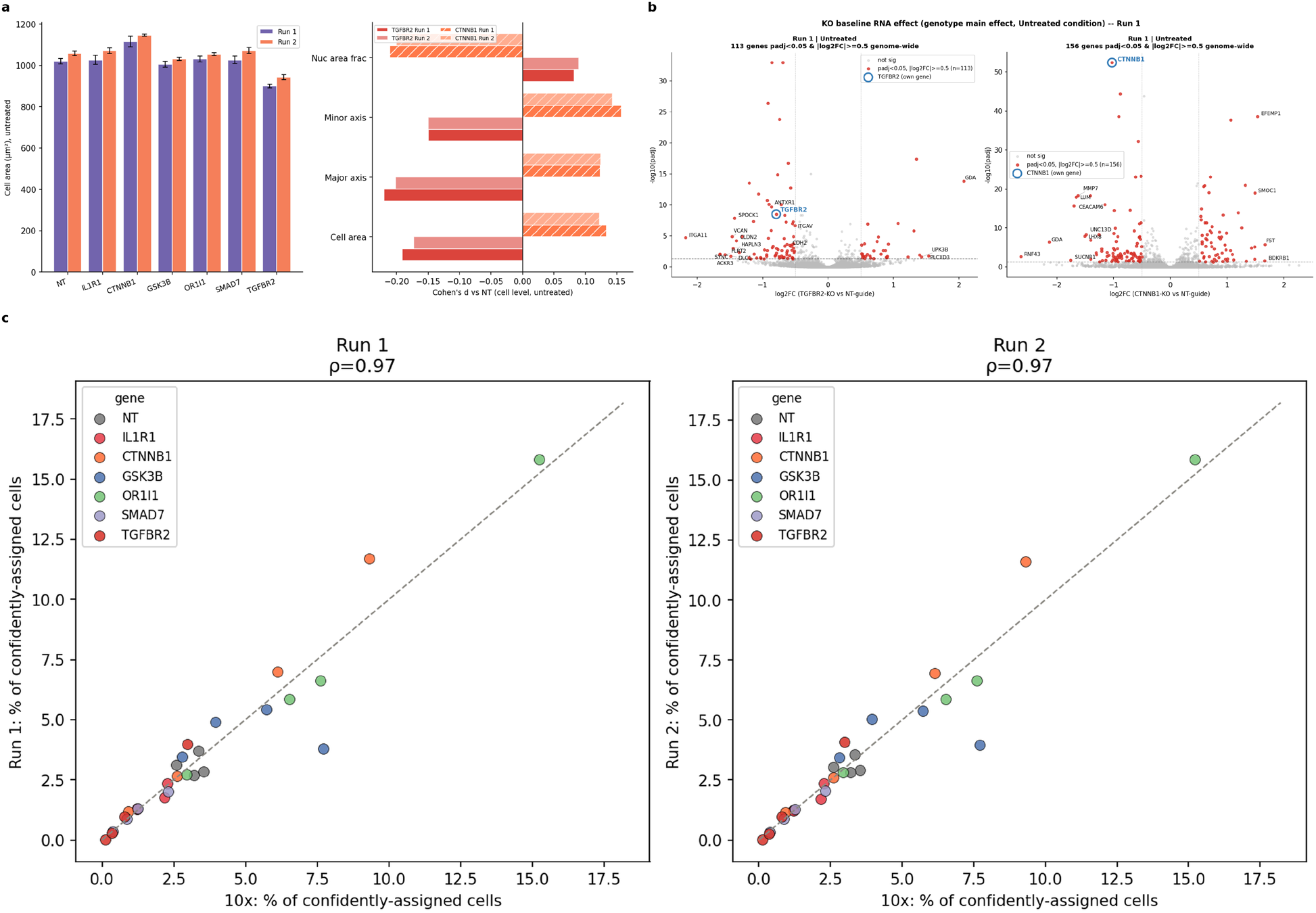
Same-cell morphology, protein and transcript in TGFBR2/CTNNB1 knockouts; guide representation against an orthogonal platform. **a**, TGFBR2 and CTNNB1 knockout: cell-area change and cell-geometry Cohen’s-d profile (area, major/minor axis, nuclear-area fraction), untreated, both runs. **b**, Baseline differential-expression volcanoes, TGFBR2 (left) and CTNNB1 (right), run 1. **c**, Guide-abundance correlation (Spearman ρ, 28 guides) against an independent 10x Genomics dataset.

To assess agreement with an orthogonal platform, we compared the guide and RNA readouts against an independently generated 10x Genomics 5′ gene expression and CRISPR guide feature barcoding dataset from the same guide library and cell line. Guide representation agreed closely (ρ = 0.97; 0.97; Fig. 3c). For TGFBR2 and CTNNB1, log2 fold changes correlated across platforms among genes significant on either platform (Spearman ρ = 0.69–0.75 across both perturbations and runs; Methods). Gene-detection saturation curves provide an additional comparison (Extended Data Fig. 10).

This cross-platform agreement supports the transcriptional readout. By measuring morphology, protein, guide identity and transcript in the same individual cells, Cytos MPS directly links genetic perturbations to multimodal cellular responses, replacing post-hoc reconciliation of separate single-modality screens.

## Methods (for online placement)

### Study design

CRISPR loss-of-function screen in A549 human lung adenocarcinoma cells stimulated with recombinant IL-1β at 15 ng/mL. Two independent biological replicate runs (run 1 and run 2) were performed on an AVITI24 instrument (Element Biosciences)^11^. Each run used a 12-well slide: four untreated wells, four wells treated with IL-1β for 15 minutes, and four for 30 minutes, all fixed concurrently at the longest timepoint. Every well contained the full pooled population of CRISPR knockout guides, so genotype is a within-well variable and each knockout-versus-control comparison is internally controlled for ligand aliquot, fixation and imaging field.

### Cell culture, transduction and fixation

A549 cells were cultured in F12K medium (Corning 10-025-CV) supplemented with 10% FBS (Gibco A56705-01), 1% penicillin/streptomycin (Gibco 15140-122), 10 µg/mL blasticidin (Gibco A1113903) and 2 µg/mL puromycin (Thermo Scientific J67236.8EQ) to maintain selection pressure. Cells were seeded in T225 flasks (Thermo Scientific 159934) at 2 × 10^6^ cells and incubated at 37 °C in a humidified 5% CO_2_ incubator, passaging every 2–3 days at 70% confluency. Bare glass slide kits (Element Biosciences 810-00012) were coated according to the user manual before seeding: washed with 0.1 M NaOH, coated with poly-L-lysine (MilliporeSigma P4707) and dried. Cells were lifted with 0.25% trypsin-EDTA (Gibco 25200-056), inactivated with culture medium, counted on a Countess 3 (Invitrogen AMQAX2000) and confirmed >90% viable. 100 µL of medium was added to each well, then 1 × 10^4^ cells per well, distributed evenly, and incubated at 37 °C overnight. IL-1β (R&D Systems 201-LB-010) was diluted to 15 ng/mL in culture medium and added by medium swap for 15 or 30 minutes before fixation; control wells received a medium swap for 30 minutes. Cells were washed three times with DPBS (Gibco 14190144), fixed in 4% formaldehyde (Fisher Scientific BP531-25) for 30 minutes, washed three times with 1× PBS (Gibco 10010049) and stored in 0.1 U/µL RiboLock (Thermo Scientific EO0381) in PBS.

### CRISPR library

Cells were transduced with a Cellecta pooled lentiviral guide RNA library targeting six genes plus non-targeting controls: NT (4 guides), CTNNB1, GSK3B, IL1R1, OR1I1, SMAD7 and TGFBR2 (4 guides each; 28 guides total), at low MOI to favor single-guide-per-cell delivery. OR1I1, an olfactory receptor with no expected phenotype in this cell type, served as an additional negative control.

### Optical pooled sequencing

Guide identity was determined in situ by optical pooled sequencing of the Cellecta B01 barcode library. Six batches of direct in situ sequencing (DISS) were performed with the Cytos RNA cartridge: one batch of 21 cycles to determine guide identity and five batches of 51 cycles to read 3′ mRNA per cell. Guide identity was determined by reverse transcription (RT) and sequencing of the gRNAs in the Cellecta barcode library: 25 µL of 2 µM Cellecta RT primer (5′-/5Phos/CAGCCGCATCTTCTTTTGCCACTTTTTCAAGTTGATAACGGACTAGCCT-3′) was spiked into the RNA RT Primer well and 25 µL of Cellecta sequencing primer (5′-GCTATGCTGTTTCCAGCATAGCTCTGAAAC-3′) into the RNA Custom Sequencing Primer well of the cartridge. 3′ mRNA counts per cell were obtained by RT and sequencing of polyA transcripts. A panel of eight ACTB and GAPDH targets was additionally spiked into the RNA RT Primer well as run controls. Per-cell guide assignments were read directly from the platform’s target-assignment fields (MostAbundantTarget, MostAbundantTargetCount, TotalCount, Confidence); none are re-derived by us. The platform defines Confidence (hereafter “assignment score”) as MostAbundantTargetCount / (TotalCount + 1) (Laplace-smoothed). Cells with assignment score strictly greater than 0.5 were retained for all guide-resolved comparisons (207,577 of 241,512; 208,507 of 239,559 RNA-QC-passing cells, 85.9%; 87.0%, matching the platform’s own reported per-run assignment rate). Guide identity was joined to RNA and imaging cells via a shared per-cell key (Well, Cell), asserted unique per run.

### RNA processing

The raw 3′ untargeted panel (54,949 features) was restricted to the 20,063 protein-coding-biotype features. Cells with mitochondrial read fraction >20% were excluded; genes detected in fewer than 10 cells (post cell-QC) were excluded, leaving 17,805; 17,780 protein-coding genes per run (Fig. 1d’s “full protein-coding panel” reference line). Extended Data Fig. 10’s “shared AVITI24 protein-coding panel” (19,288 genes) is the subset of the 10x Genomics dataset’s own gene panel that also falls within AVITI24’s protein-coding gene set, used as the detection denominator for the 10x curve specifically; it is a different quantity from the per-run AVITI “full protein-coding panel” counts above, not a third independent estimate of the same thing. Counts are raw, direct-sequencing molecule counts (no PCR amplification step, no UMI correction). No bulk normalization was applied; pseudobulk differential expression used PyDESeq2^12^ (two-sided Wald test) with its own size-factor normalization on raw integer counts, well × class aggregation (≥10 qualifying cells/well), Benjamini-Hochberg FDR < 0.05 genome-wide. The log2 fold-change cutoff is |log2FC| ≥ 0.5 for knockout baseline comparisons (Fig. 3b; Extended Data Figs. 8, 9) and |log2FC| > 0.5 for the IL-1β induction comparison (Fig. 1e); volcano plots show these tests.

### Protein panel and localization scoring

Protein is measured by automated immunofluorescence; the platform supports up to 20 protein measurements per run, independent of isotype controls and Cell Painting markers. This study used 18 antibody reagents: 15 protein-directed readouts, one biotin-conjugation control, and two isotype controls (see Supplementary Table 1 for the full list: clone, vendor, catalog number), each scored as 10 compartment- and intensity-resolved per-cell features (whole-cell, nuclear, cytoplasmic, threshold-exceeding fraction, and nuclear:whole-cell localization ratio). The 15 protein-directed readouts plus the conjugation control (16 non-isotype antibodies) each contribute 10 features, yielding 160 base features (150 protein-directed and 10 from the conjugation control). The panel spans an NF-κB module (NF-κB p65/RelA, phospho-NF-κB sites), a MAPK module (phospho-p38, phospho-HSP27, phospho-ERK1/2 and total ERK1/2, measured with separate antibodies), and structural/organelle markers. Two anti-p65 reagents are used: A19653BC (feature prefix NFkB_BC), a site-specific antibody, is the primary NF-κB readout used throughout; A19653BCR (feature prefix NFkB_BCR) is a random-biotin conjugate of the same clone and serves as a conjugation control, not an independent protein target. We refer to the NFkB_BC readout as “NF-κB p65” or “NF-κB nuclear fraction” in prose and figures; exact platform feature identifiers (NFkB_BC, NFkB_BCR) are retained in Methods and in feature-level figures (Extended Data Fig. 7). Per-cell nuclear-to-whole-cell ratio (platform field NuclearRatio) is a threshold-based localization score, undefined when no pixel in a cell clears that channel’s detection threshold.

### Single-cell NF-κB/phospho coupling

Non-targeting-guide and IL1R1-knockout cells (assignment score >0.5) were binned into 10 equal-count deciles by NFkB_BC nuclear:whole-cell ratio, separately per run, condition (untreated, 15 min, 30 min) and guide; decile-mean phospho-p38 and phospho-HSP27 whole-cell intensity were plotted against decile-mean NFkB_BC, and Spearman’s ρ was computed on the underlying per-cell values within each run/condition/guide/marker combination. To check whether cell size confounds this relationship, we additionally computed an area-adjusted Spearman correlation controlling for per-cell area (`Area_um2`), regressing area out of the NFkB_BC and phospho values via a linear fit within each run/condition/guide group before recomputing ρ on the residuals.

### Same-cell NF-κB/RNA decile test

Using the same per-cell NFkB_BC nuclear-fraction deciles as above, non-targeting-guide cells were joined to their matched RNA profile via the shared per-cell key (95.3%; 95.6% of RNA-QC-passing cells had a matching imaging cell, both runs), and pseudobulk CPM for NFKBIA and DUSP1 was computed within each decile (summed raw counts / summed library size × 10^6^, Laplace-smoothed by one read to avoid instability at near-zero counts). Effect size is reported as the decile-10:decile-1 pseudobulk CPM ratio, with a 95% confidence interval from resampling cells with replacement independently within each of the two extreme deciles (2,000 resamples). To assess reproducibility, phospho correlations (including area adjustment) and RNA ratios were also calculated separately within each well in stimulated non-targeting cells. RNA deciles were redefined within wells; estimates were summarized as mean ± s.e.m. across four wells separately for each run. Cell-bootstrap intervals quantify pooled-cell uncertainty, not between-well variability. Per-cell detected/undetected splits of individual transcripts were not used for the same-cell test because, at this depth, such splits are confounded with per-cell total counts; deciles were therefore defined on the imaging readout and RNA evaluated as pseudobulk with depth and background-gene controls (Extended Data Fig. 5).

### Confound check for the same-cell RNA decile test

To test whether NFKBIA/DUSP1’s rise (above) reflects NF-κB-target specificity rather than a generic per-cell-activity confound (e.g. cell size, cell-cycle phase, or overall transcriptional output rising together with nuclear NF-κB), we compared their decile-10:decile-1 ratio against a genome-wide background of stably-expressed genes rather than a small hand-picked panel. Per run, candidate background genes were detected above a false-positive-calibrated CPM threshold (CPM ≥1.14; 1.01), calibrated on the cells used in this test to 0% FPR against olfactory-receptor negative controls as for Extended Data Fig. 10, excluding genes on a curated NF-κB-target/immediate-early/AP-1 list, mitochondrial genes, and NFKBIA/DUSP1 themselves (11,985; 12,094 genes; this calibration uses a different cell population, condition and CPM threshold than the headline detected-gene counts in Fig. 1d, so the two are not directly comparable). Because a CPM-based detection call can still reflect as little as one read in a single well’s extreme-decile bin — too little to support a reliable per-well ratio — genes were further required to contribute at least 20 raw reads to both the decile-1 and decile-10 pseudobulk sum in every well, leaving 1,621; 1,673 background genes per run. For each background gene and for NFKBIA/DUSP1, one decile-10:decile-1 pseudobulk CPM ratio was computed per well (matching the well-level replicate unit used elsewhere in this manuscript, n=4 wells/condition) and averaged across wells, with s.e.m. across wells; NFKBIA/DUSP1’s percentile rank was then read off the resulting background distribution. NFKBIA/DUSP1 exceeded 91.1–99.9% of the background distribution in both runs (Extended Data Fig. 5), arguing against a generic per-cell-activity confound as the driver of the Fig. 2d rise.

### Imaging

Cells require a detected nucleus and fall within the per-well 1st–99th percentile of cell area. Morphology geometry (area, major/minor axis, nuclear area fraction) is taken directly from platform-provided per-cell geometry from an automated segmentation model (cellpose3). Cell Painting^13^ utilizes commercially available reagents including dye-labelled WGA, phalloidin and RedDot2. Modalities are registered to one another via the shared (Well, Cell) key assigned at segmentation, not by independent image alignment. The CellPaint panel (actin, cell membrane, mitochondria, nucleus; `target_type==‘CellPaint’, feature_class==‘base’`, 40 features) and the protein/IF panel (160 features) were each tested per feature for a knockout-vs-non-targeting difference (Cohen’s d, Mann-Whitney U, Benjamini-Hochberg FDR across the panel), Untreated condition, assignment score >0.5 cells only. Knockout cell-area change was tested by paired per-well t-test (n = 4 wells per run).

### Cross-platform comparison

An independently generated 10x Genomics 5′ gene expression and CRISPR guide feature barcoding dataset (same pooled guide library, A549 cells) was compared against both AVITI24 runs. Relative guide abundance was compared by Spearman correlation across all 28 guides, computing each guide’s percentage relative to its own platform’s assigned population (AVITI24: score >0.5; 10x: cells with exactly one Cellecta-panel guide in the call list), so both platforms’ percentages sum to 100% and are directly comparable. Differential expression in the 10x data used per-cell, per-gene Mann-Whitney U tests (target guide vs. non-targeting) with genome-wide Benjamini-Hochberg correction; the AVITI24 comparison used pseudobulk differential expression instead, using four wells per condition within each run; the 10x dataset lacked replicate structure. Gene-detection sets on each platform were defined by pooled-CPM thresholds calibrated against a shared negative-control gene panel (olfactory receptor genes) to a target false-positive rate of 0%. Gene-detection saturation curves (Fig. 1d; Extended Data Fig. 10) subsample cells from a fixed pool and report mean ± s.d. across resampling replicates. 10x counts are UMI-corrected following standard 10x Genomics processing; AVITI24 counts are raw, uncorrected molecule counts. For each perturbation and run, Spearman correlation of log2 fold changes was calculated among genes significant on either platform. This selected-set comparison assesses transcriptional response concordance; differing designs and tests preclude interpreting hit counts as relative sensitivity or specificity.

### Statistics and reproducibility

No statistical method was used to predetermine sample size; sample sizes reflect the full pooled-guide cell population passing QC in each well. No data were excluded except by the QC criteria specified above. Investigators were not blinded, as all comparisons were performed computationally across the full cell population within each well rather than by manual scoring. Effect sizes are reported as cell-level Cohen’s d (sample SD, ddof=1) throughout (Fig. 3a; Extended Data Figs. 2, 7). Cell-level two-sided Mann-Whitney U tests with Benjamini-Hochberg correction are used for single-cell distribution comparisons (Extended Data Fig. 2) and for the CellPaint and protein-panel feature screens (Extended Data Fig. 7). Well-level tests (n=4 wells/condition) are used for the IL1R1 attenuation P values in the main text and the knockout cell-area change (paired t-test). The confound-check background distribution (Extended Data Fig. 5) and other well-level summaries in figures are reported as mean ± s.e.m. across the n=4 replicate wells per run, a variability summary rather than a hypothesis test. Two-sided tests throughout; multiple-testing correction as specified per comparison (genome-wide BH-FDR unless otherwise noted). n = 2 independent biological replicate runs throughout.

## Data availability

Spatial data objects, including images and per-cell measurements for runs 1 and 2, will be deposited in the BioImage Archive. The 10x Genomics comparison dataset will be deposited in GEO/SRA. Accession numbers will be added when available, and the datasets will be publicly released no later than journal publication.

## Code availability

Analysis code used to generate the figures and statistics will be made publicly available on GitHub before journal publication, with a versioned release archived in Zenodo.

## Acknowledgements

We thank Joseph Puglisi for valuable feedback on earlier versions of the manuscript.

## Author contributions

T.L., J.R., B.S., A.B. performed experiments; D.H., M.K., J.Z., X.Q. developed chemistry/instrumentation; T.L., S.K. designed the study; S.K., T.L. and A.S. analyzed data; S.K. wrote the manuscript with input from all authors.

## Competing interests

All authors are employees of Element Biosciences and may hold stock or stock options in the company. Element Biosciences develops and sells the AVITI24 instrument and the Cytos MPS workflow described here.

**Supplementary Table 1.**
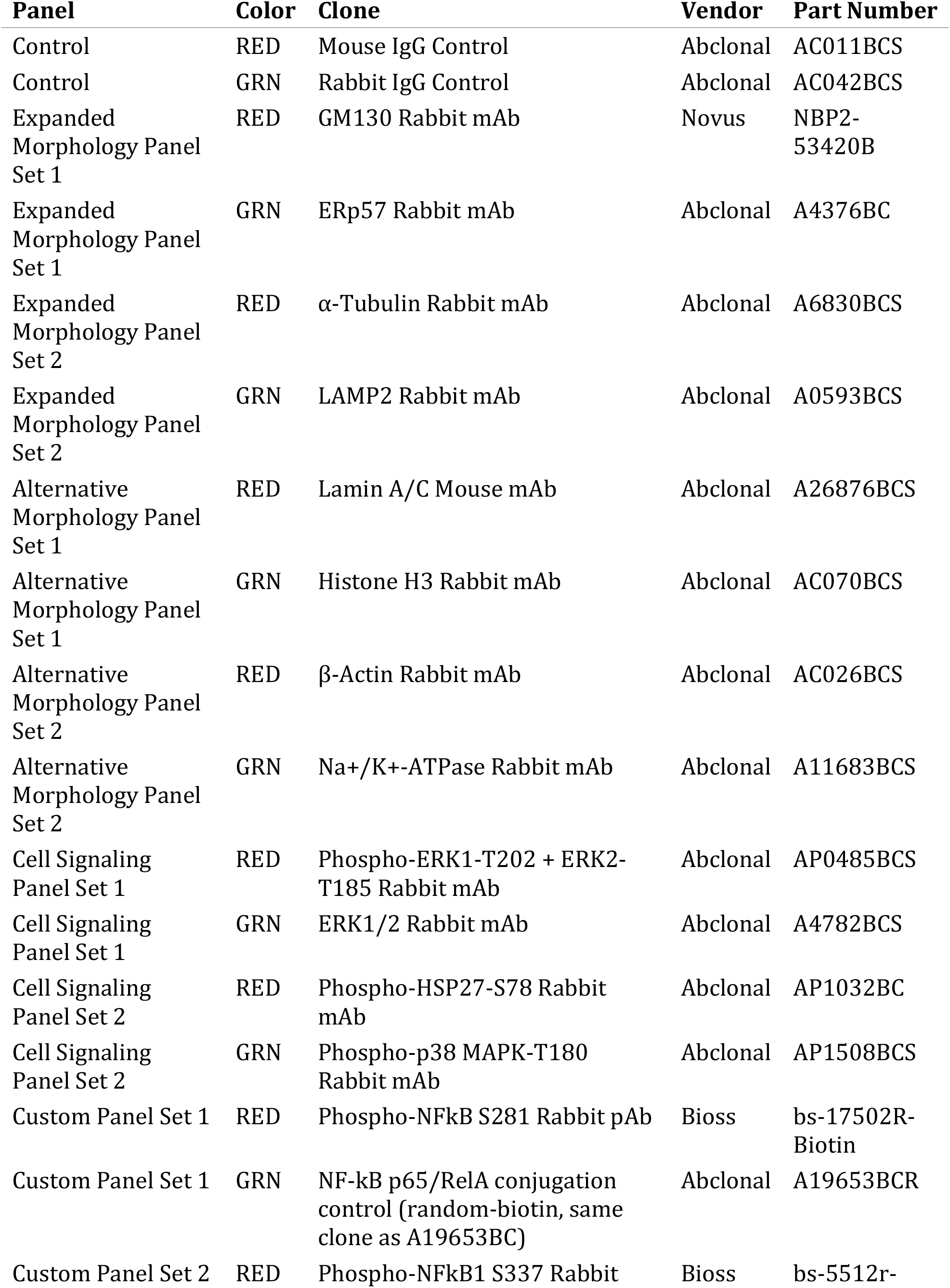
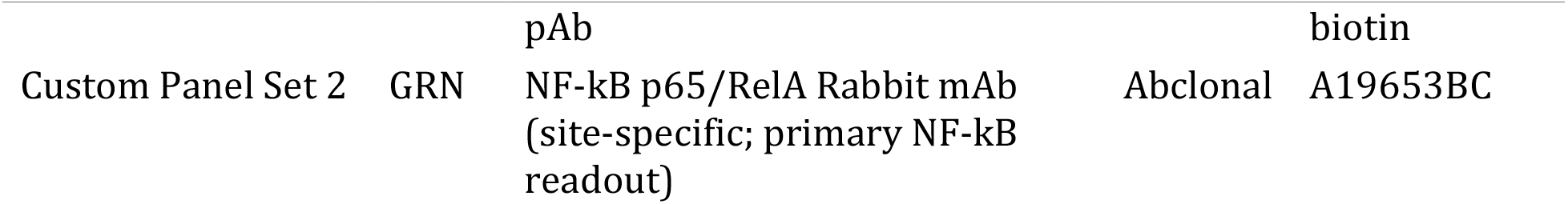
Antibody panel. Panel set, imaging colour, clone, vendor and part number for the 15 protein-directed readouts, one biotin-conjugation control (A19653BCR) and two isotype controls.

| **Panel** | **Color** | **Clone** | **Vendor** | **Part Number** |
| --- | --- | --- | --- | --- |
| Control | RED | Mouse IgG Control | Abclonal | AC011BCS |
| Control | GRN | Rabbit IgG Control | Abclonal | AC042BCS |
| Expanded Morphology Panel Set 1 | RED | GM130 Rabbit mAb | Novus | NBP2-53420B |
| Expanded Morphology Panel Set 1 | GRN | ERp57 Rabbit mAb | Abclonal | A4376BC |
| Expanded Morphology Panel Set 2 | RED | α-Tubulin Rabbit mAb | Abclonal | A6830BCS |
| Expanded Morphology Panel Set 2 | GRN | LAMP2 Rabbit mAb | Abclonal | A0593BCS |
| Alternative Morphology Panel Set 1 | RED | Lamin A/C Mouse mAb | Abclonal | A26876BCS |
| Alternative Morphology Panel Set 1 | GRN | Histone H3 Rabbit mAb | Abclonal | AC070BCS |
| Alternative Morphology Panel Set 2 | RED | β-Actin Rabbit mAb | Abclonal | AC026BCS |
| Alternative Morphology Panel Set 2 | GRN | Na<sup>+</sup>/K<sup>+</sup>-ATPase Rabbit mAb | Abclonal | A11683BCS |
| Cell Signaling Panel Set 1 | RED | Phospho-ERK1-T202 + ERK2-T185 Rabbit mAb | Abclonal | AP0485BCS |
| Cell Signaling Panel Set 1 | GRN | ERK1/2 Rabbit mAb | Abclonal | A4782BCS |
| Cell Signaling Panel Set 2 | RED | Phospho-HSP27-S78 Rabbit mAb | Abclonal | AP1032BC |
| Cell Signaling Panel Set 2 | GRN | Phospho-p38 MAPK-T180 Rabbit mAb | Abclonal | AP1508BCS |
| Custom Panel Set 1 | RED | Phospho-NFkB S281 Rabbit pAb | Bioss | bs-17502R-Biotin |
| Custom Panel Set 1 | GRN | NF-kB p65/RelA conjugation control (random-biotin, same clone as A19653BC) | Abclonal | A19653BCR |
| Custom Panel Set 2 | RED | Phospho-NFkB1 S337 Rabbit | Bioss | bs-5512r- |

| Panel | Color | Clone | Vendor | Part Number |
| --- | --- | --- | --- | --- |
|  |  | pAb |  | biotin |
| Custom Panel Set 2 | GRN | NF-kB p65/RelA Rabbit mAb<br>(site-specific; primary NF-kB readout) | Abclonal | A19653BC |

## Extended Data Figure legends

**Extended Data Fig 1.**
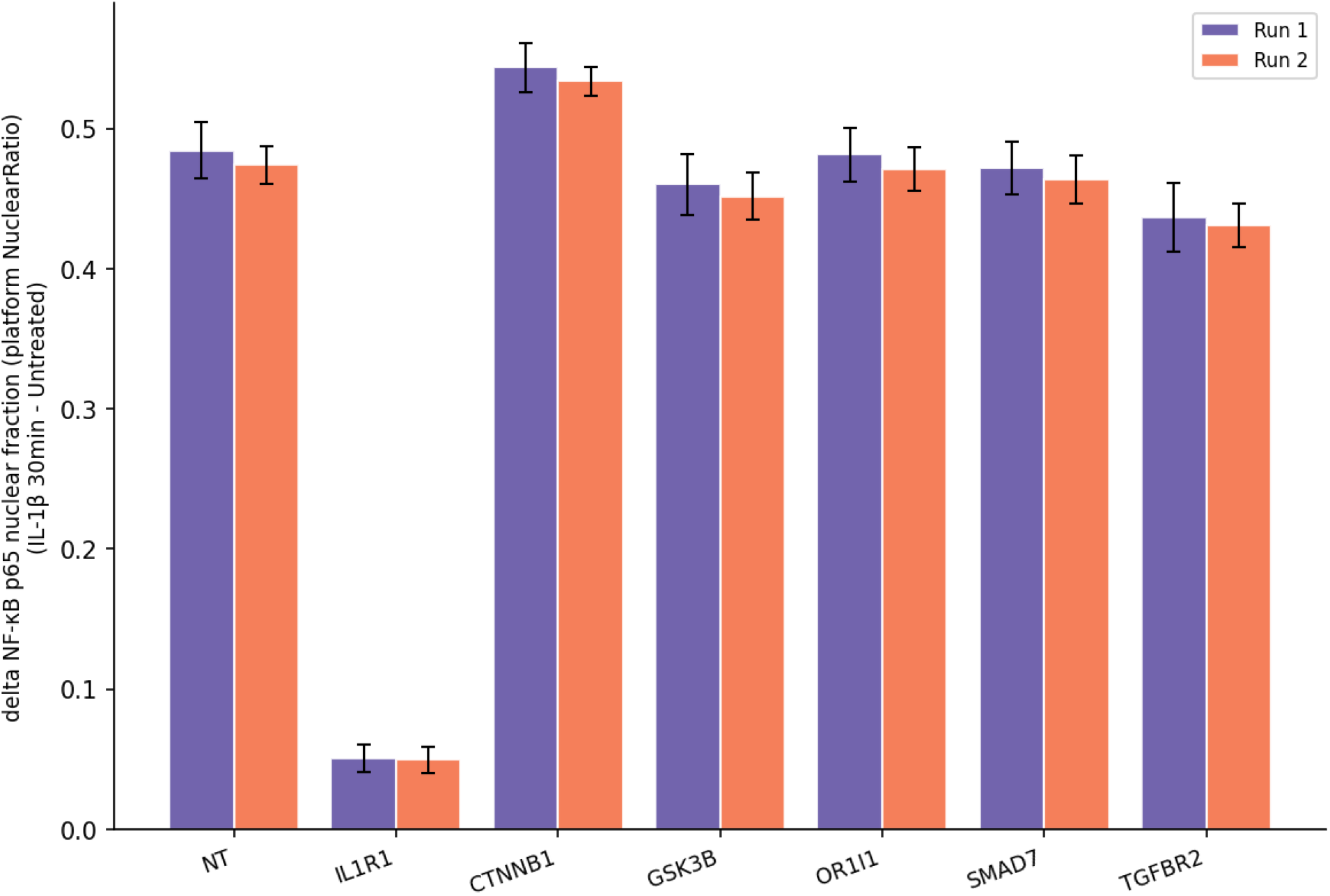
Per-knockout NF-κB translocation deltas. Per-well mean NF-κB nuclear-fraction shift (30-min stimulated minus untreated), all 6 knockout genes plus non-targeting, both runs.

**Extended Data Fig 2.**
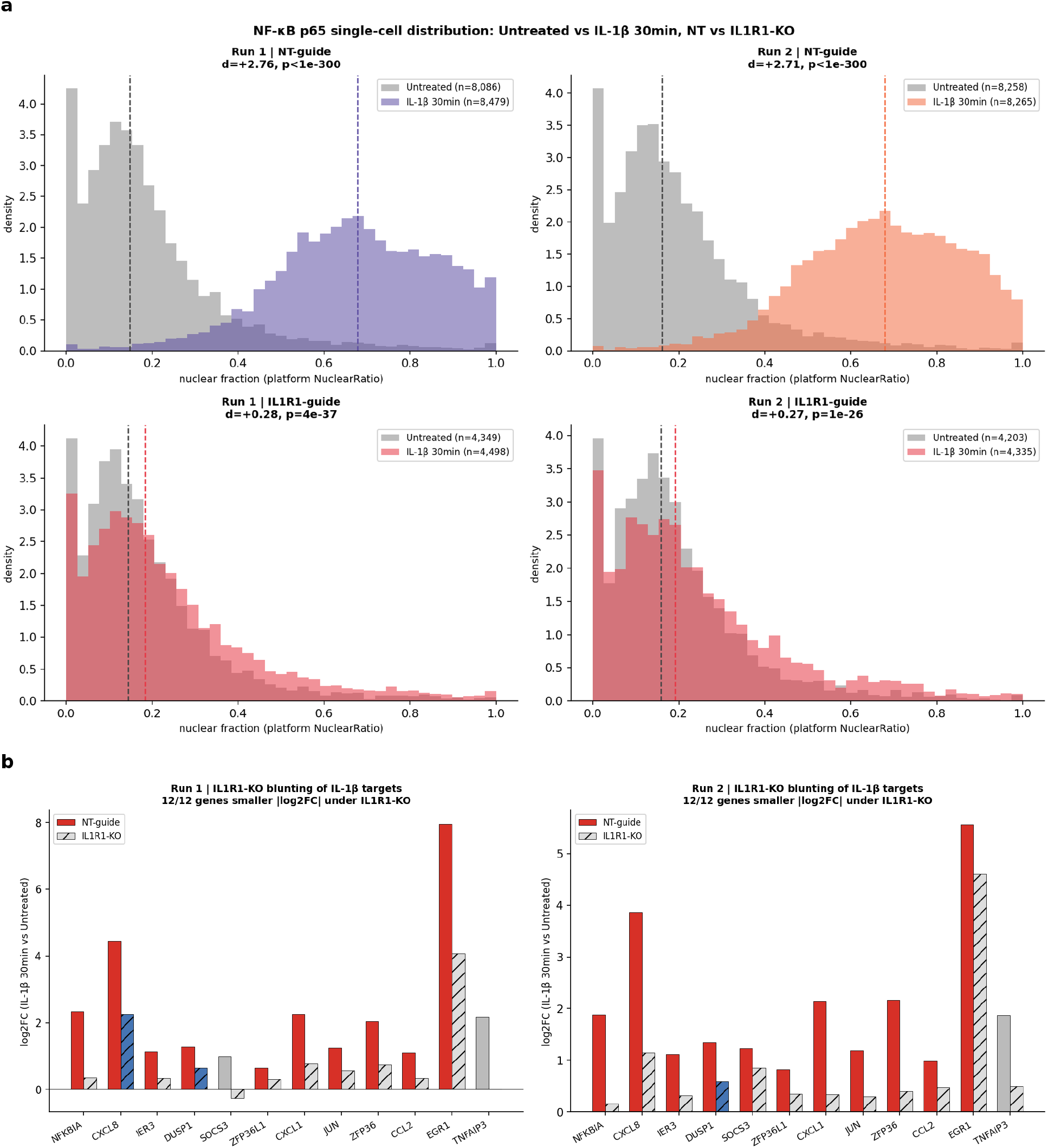
IL1R1 knockout blunts the IL-1β response at protein and transcript level. **a**, Single-cell NF-κB nuclear-fraction distributions, untreated (grey) vs. 30-min stimulated (colored), non-targeting vs. IL1R1-KO; dashed vertical lines mark condition medians; Cohen’s d and two-sided Mann-Whitney U P annotated per panel. **b**, IL-1β 30-min vs. untreated fold-change for the 12 IL-1β-induced genes in non-targeting vs. IL1R1-KO cells, both runs, coloured by significance in non-targeting (red), IL1R1-KO (blue) or neither (grey).

**Extended Data Fig 3.**
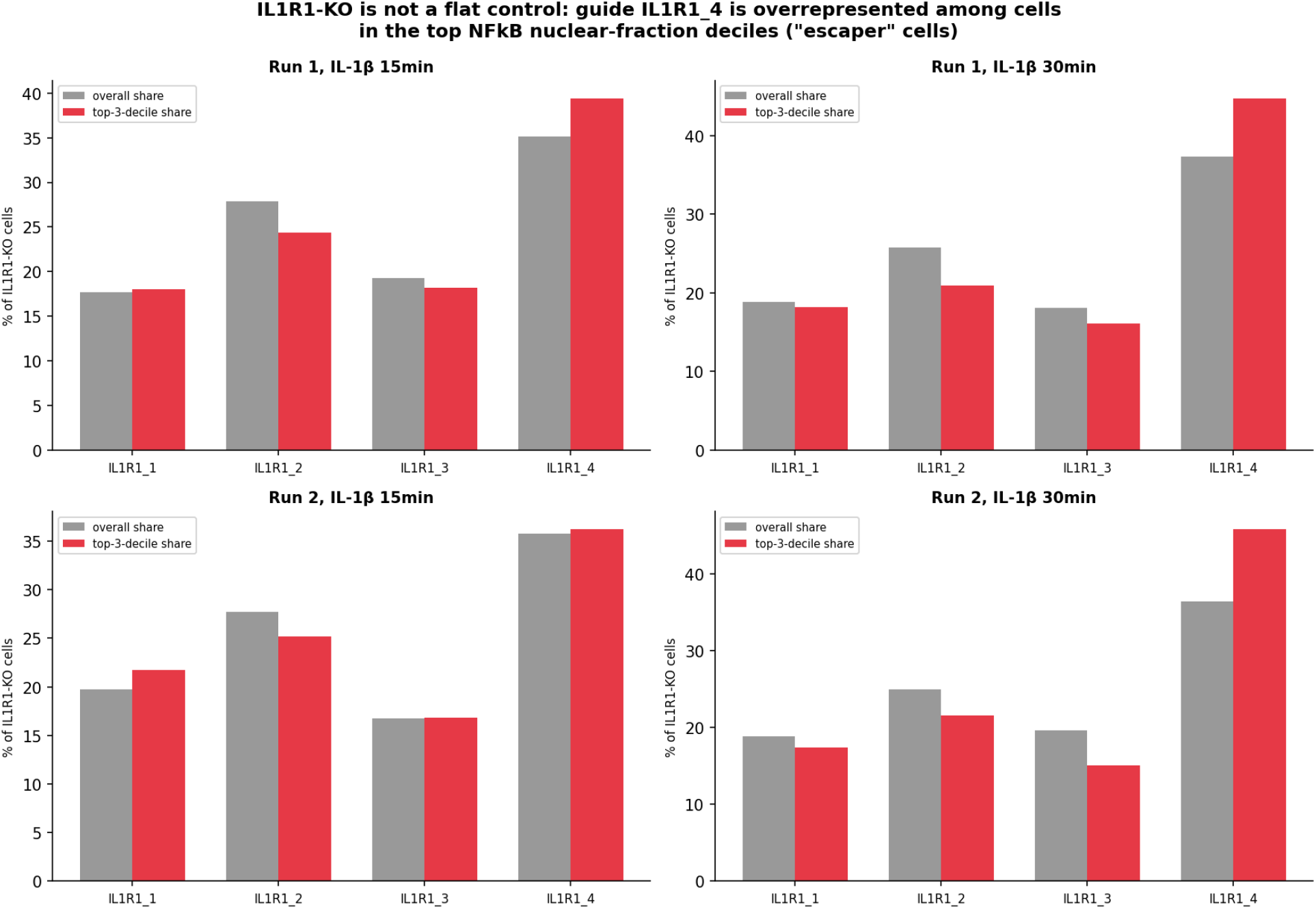
Guide-level breakdown of IL1R1-knockout cells with elevated nuclear NF-κB. Guide share among IL1R1-knockout cells, overall vs. restricted to the top-3 NF-κB-nucratio deciles, 15-min and 30-min, both runs; one guide (IL1R1_4) is overrepresented among cells that still show elevated nuclear NF-κB despite knockout.

**Extended Data Fig 4.**
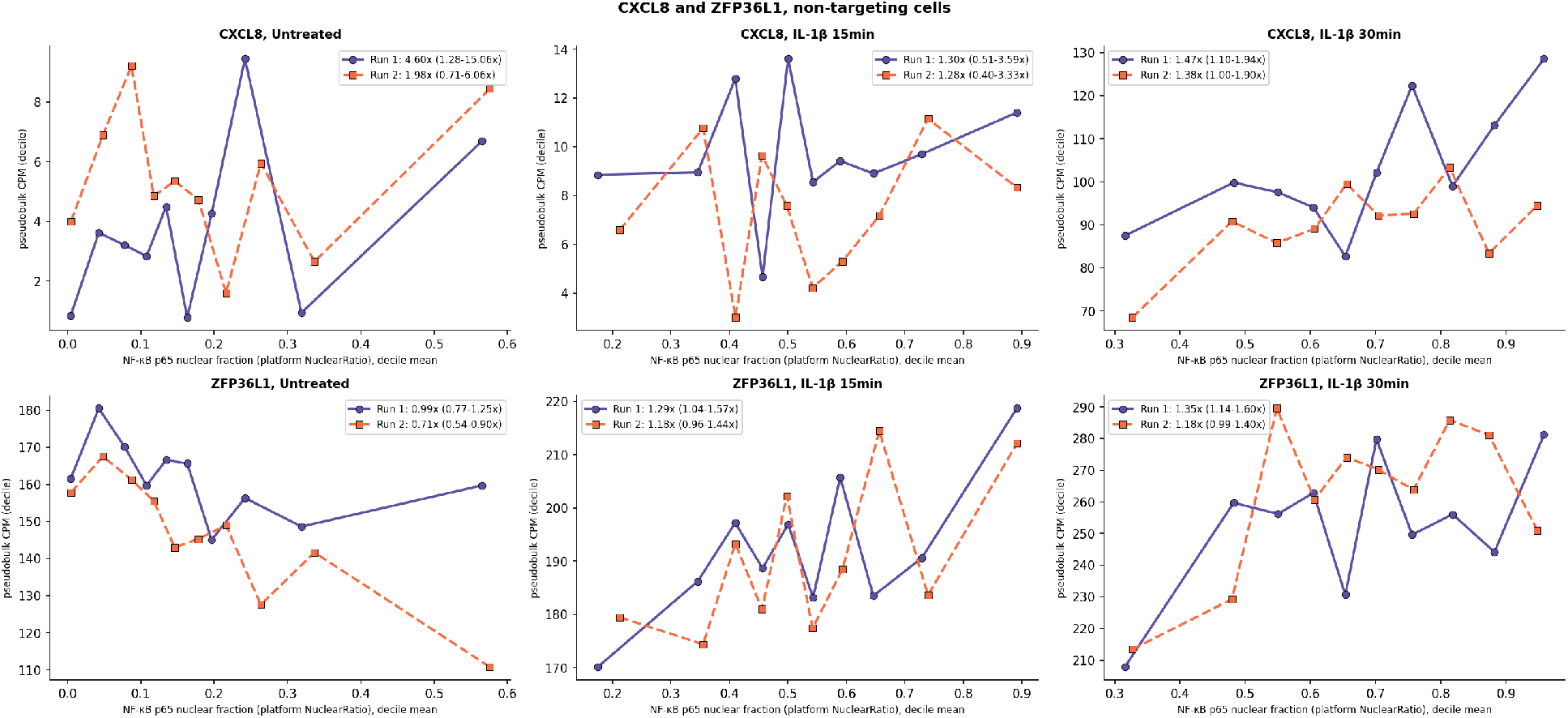
CXCL8 and ZFP36L1 same-cell RNA-decile analysis. CXCL8 and ZFP36L1, same layout as Fig. 2d, non-targeting cells only; effect sizes are smaller and less consistent across runs than for NFKBIA/DUSP1.

**Extended Data Fig 5.**
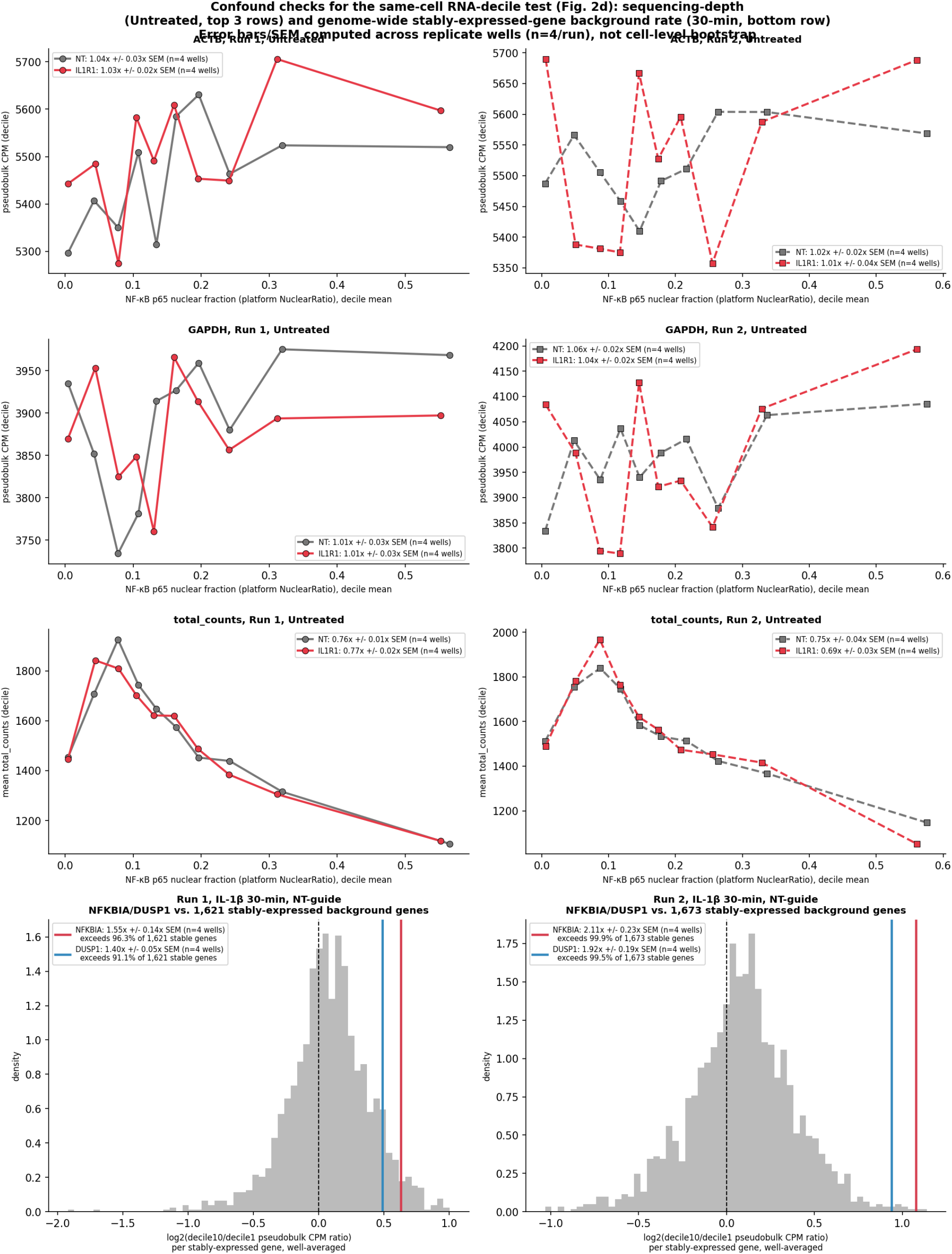
Confound checks for the same-cell RNA decile test. Sequencing-depth check (top three rows): ACTB/GAPDH pseudobulk CPM and mean total_counts vs. NF-κB-nucratio decile, Untreated cells only, both genotypes/runs — housekeeping genes stay flat across deciles even though raw per-cell library size declines, arguing against sequencing depth as the driver of the baseline DUSP1 decline noted in Fig. 2d. Genome-wide stable-gene null (bottom row): distribution of well-averaged decile-10:decile-1 pseudobulk CPM ratios across 1,621; 1,673 stably-expressed background genes per run (grey; detected above a false-positive-calibrated CPM threshold (Methods), further restricted to genes with ≥20 reads in both extreme-decile bins in every well) versus NFKBIA (red) and DUSP1 (blue) at 30 minutes, non-targeting cells — NFKBIA/DUSP1 exceed 91.1–99.9% of the background distribution in both runs. Error bars/s.e.m. throughout this figure are computed across the n=4 replicate wells per run, not by cell-level bootstrap.

**Extended Data Fig 6.**
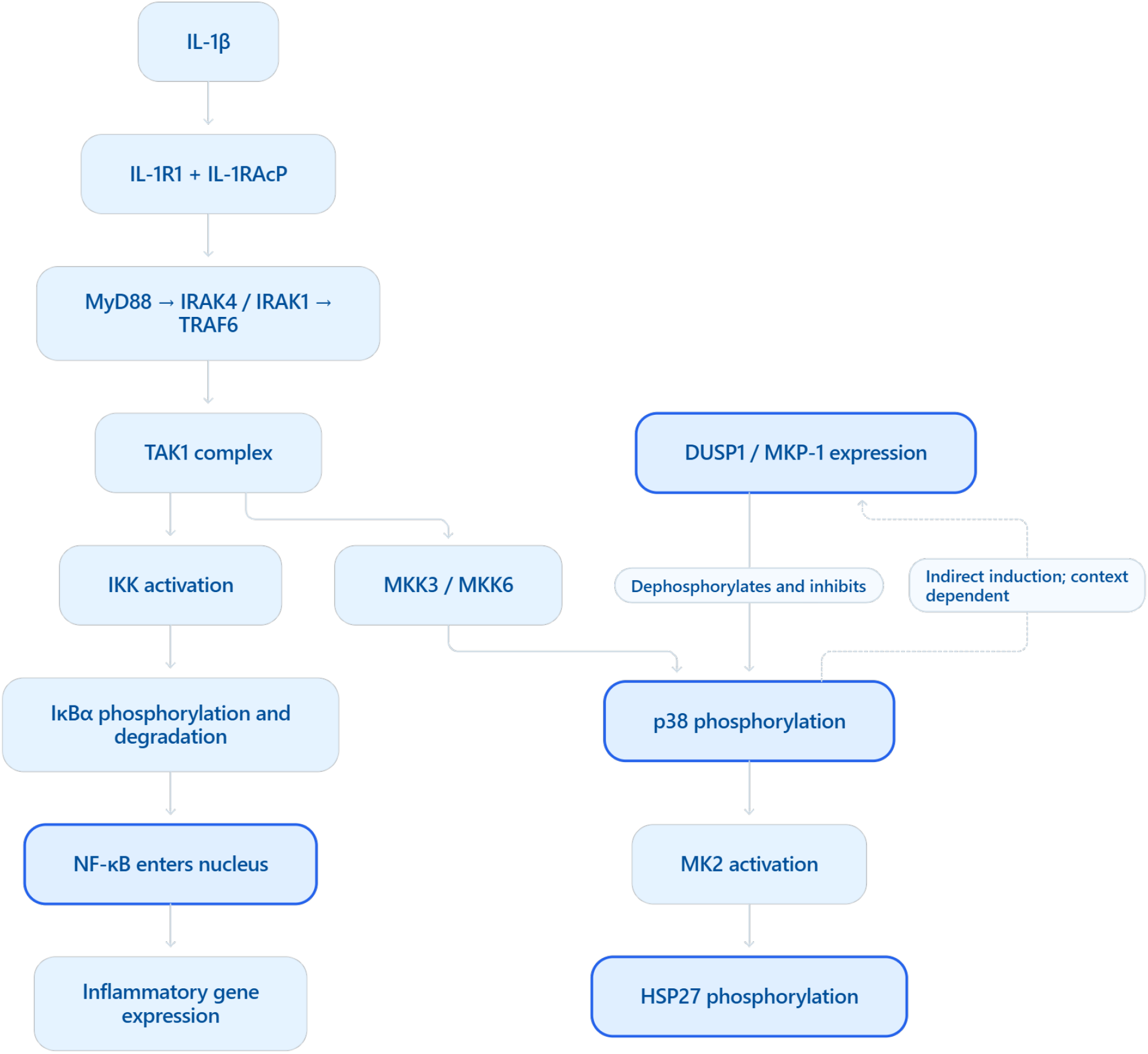
IL-1β/NF-κB and MAPK pathway schematic. Canonical IL-1β/NF-κB and MAPK pathway biology from the literature, shown for orientation; no edge in this diagram was tested in this study.

**Extended Data Fig 7.**
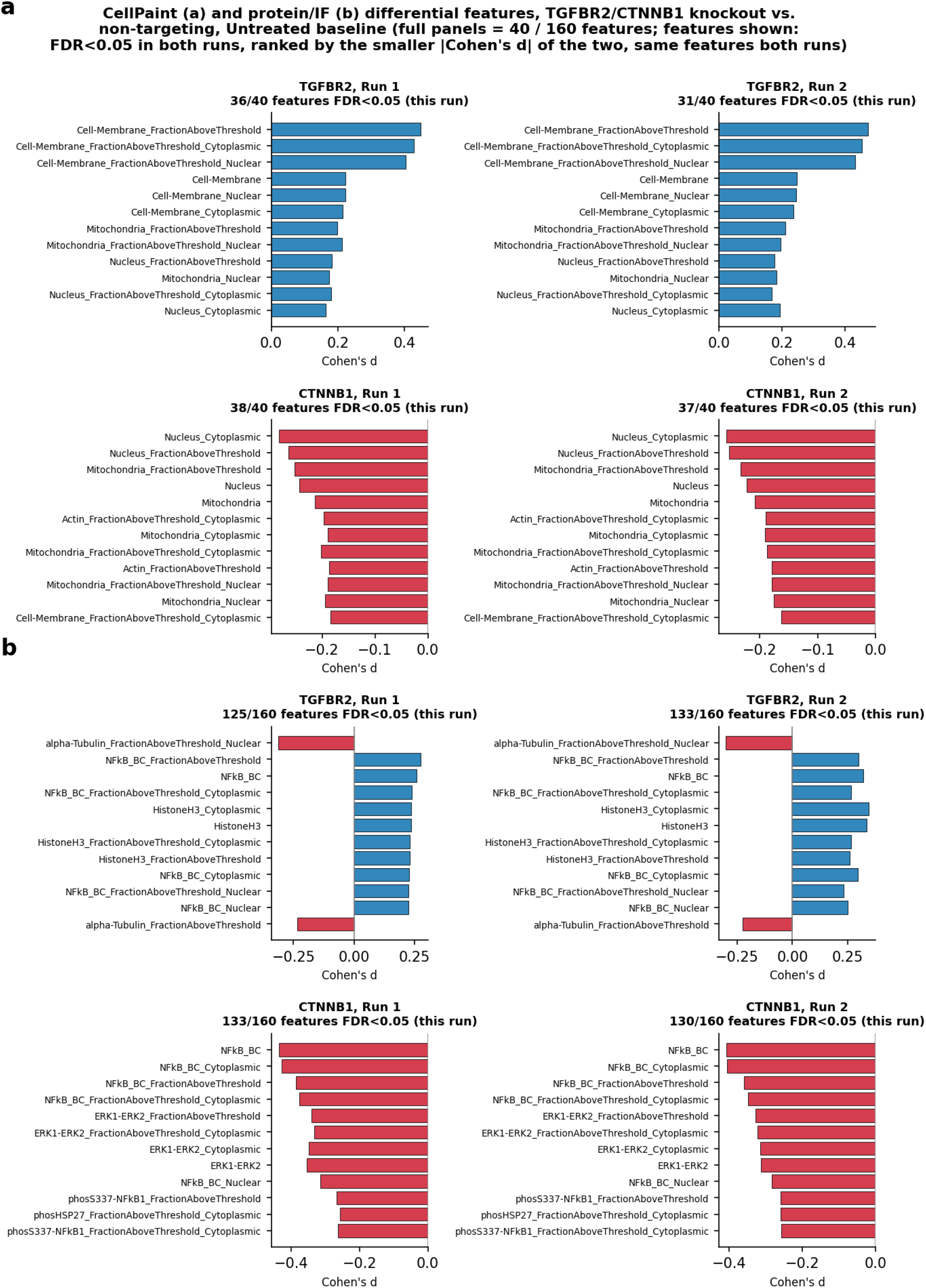
CellPaint and protein-panel differential features. TGFBR2 and CTNNB1 vs. non-targeting, Untreated baseline, both runs. **a**, CellPaint Cohen’s-d profile (40 features), led by cell-membrane signal (d = +0.45; +0.47 for TGFBR2). **b**, Protein/IF panel (full panel = 160 features; 125; 133 and 133; 130 significant, FDR < 0.05). Features shown in **b** are those significant in both runs, the same set in both panels, ranked by the smaller |Cohen’s d| of the two runs. The BCR conjugation control (random-biotin conjugate of the same p65 clone; Methods) showed little IL-1β-induced translocation (d = +0.13; +0.11, vs. +2.76; +2.71 for NF-κB p65 in non-targeting cells), indicating loss of most specific binding. Its baseline knockout shifts in **b** were smaller than NF-κB p65’s in both genotypes and runs (TGFBR2: d = +0.11; +0.16 vs. +0.26; +0.32; CTNNB1: d = −0.20; −0.17 vs. −0.44; −0.41) and shrank more under area adjustment (21–29% vs. 4–16% for NF-κB p65), supporting a p65-specific component beyond generic cell-size confounding.

**Extended Data Fig 8.**
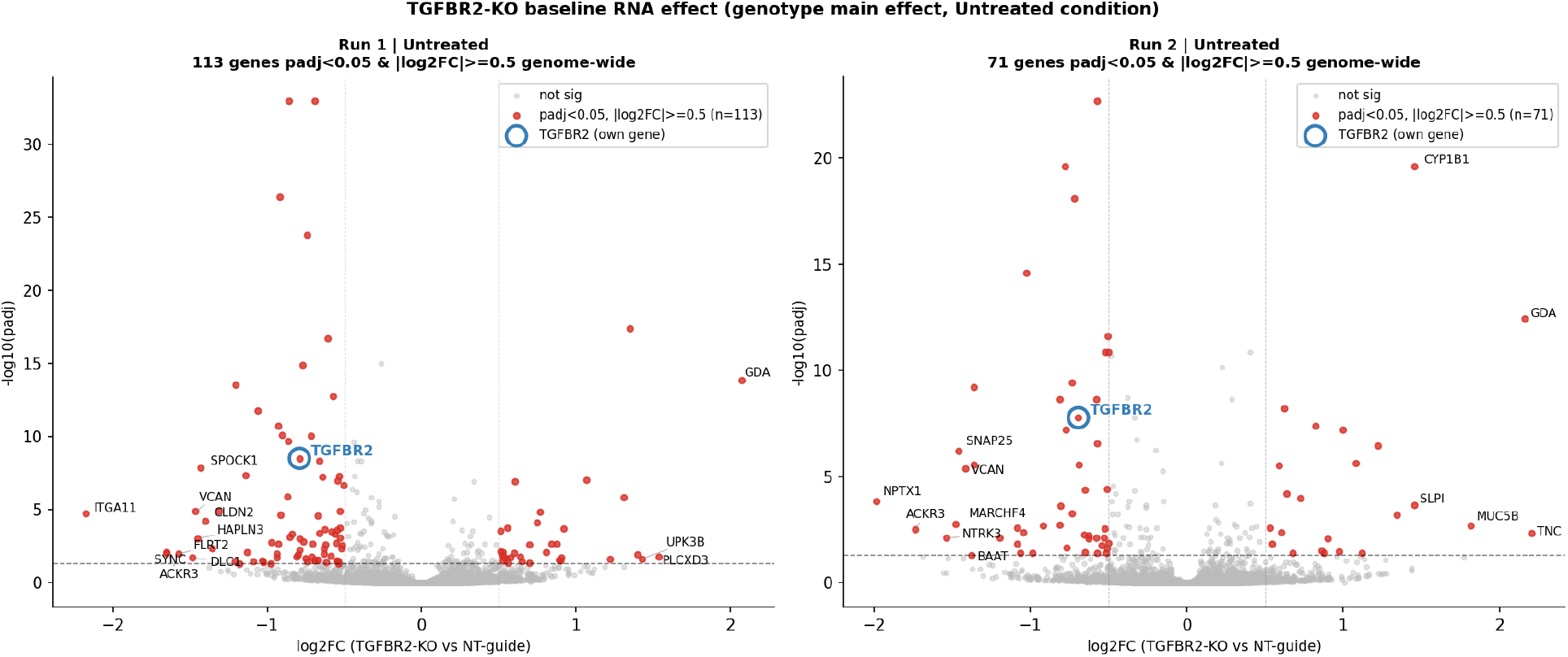
TGFBR2 baseline volcano, both runs. Both-run replicate version of Fig. 3b’s TGFBR2 panel (run 1 and run 2 shown separately).

**Extended Data Fig 9.**
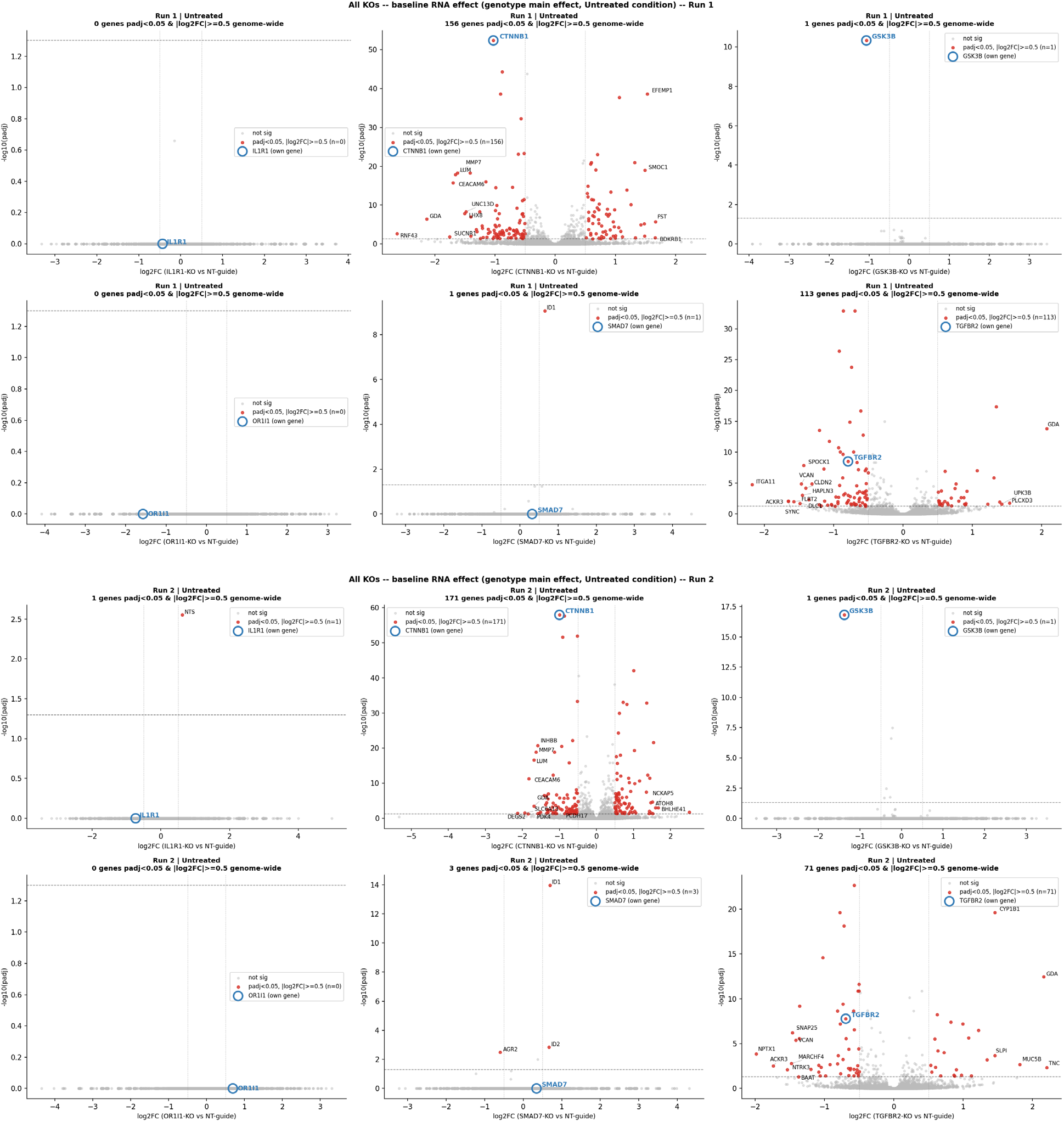
All-knockout baseline differential expression. Baseline (Untreated) volcano plots, all 6 knockout genes vs. non-targeting, (**a**) run 1 and (**b**) run 2.

**Extended Data Fig 10.**
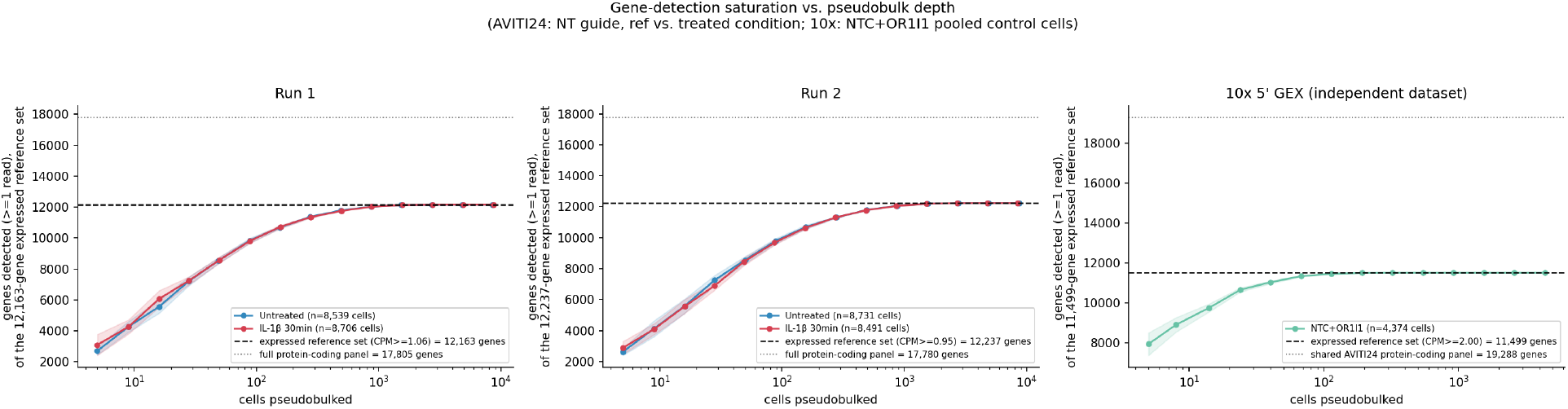
Gene-detection sensitivity vs. an orthogonal platform. Gene-detection saturation curves (run 1, run 2 and an independent 10x Genomics dataset); expressed-gene sets calibrated to 0% false-positive rate against a negative-control (olfactory-receptor) gene panel on both platforms.

## Notes

### Summary of Updates

Added author (X. Qi). Renamed platform to Cytos MPS. Clarified antibody panel composition and added analysis of the conjugation-control antibody (Extended Data Fig. 7). Reframed TGFBR2/CTNNB1 morphology results around opposite cell-geometry phenotypes and added a CellPaint feature panel (Extended Data Fig. 7a). Expanded Methods (gene-panel denominators, guide-assignment scoring, statistical tests). Renumbered Extended Data figures in citation order, corrected figure labels, and completed references. No change to data or conclusions.

## References

1. Feldman, D. et al. Optical Pooled Screens in Human Cells. Cell 179, 787–799.e17 (2019).

2. Sivanandan, S. et al. A pooled Cell Painting CRISPR screening platform enables de novo inference of gene function by self-supervised deep learning. Nat. Commun. 17, 77 (2026).

3. Dixit, A. et al. Perturb-Seq: Dissecting Molecular Circuits with Scalable Single-Cell RNA Profiling of Pooled Genetic Screens. Cell 167, 1853–1866.e17 (2016).

4. Binan, L. et al. Simultaneous CRISPR screening and spatial transcriptomics reveal intracellular, intercellular, and functional transcriptional circuits. Cell 188, 2141–2158.e18 (2025).

5. Dhainaut, M. et al. Spatial CRISPR genomics identifies regulators of the tumor microenvironment. Cell 185, 1223–1239 (2022).

6. Honigfort, D. et al. Direct In-Sample Sequencing of the 3′ Transcriptome Expands the Capabilities of Optical Pooled Screens. bioRxiv (2025). doi:10.1101/2025.10.11.681797

7. Weber, A., Wasiliew, P. & Kracht, M. Interleukin-1 (IL-1) pathway. Sci. Signal. 3, cm1 (2010).

8. Liu, Y., Shepherd, E. G. & Nelin, L. D. MAPK phosphatases — regulating the immune response. Nat. Rev. Immunol. 7, 202–212 (2007).

9. Freudlsperger, C. et al. TGF-β and NF-κB signal pathway cross-talk is mediated through TAK1 and SMAD7 in a subset of head and neck cancers. Oncogene 32, 1549–1559 (2013).

10. Ma, B. & Hottiger, M.O. Crosstalk between Wnt/β-Catenin and NF-κB Signaling Pathway during Inflammation. Front. Immunol. 7, 378 (2016).

11. Arslan, S. et al. Sequencing by avidity enables high accuracy with low reagent consumption. Nat. Biotechnol. 42, 132–138 (2024).

12. Muzellec, B., Teleńczuk, M., Cabeli, V. & Andreux, M. PyDESeq2: a python package for bulk RNA-seq differential expression analysis. Bioinformatics 39, btad547 (2023).

13. Bray, M.-A. et al. Cell Painting, a high-content image-based assay for morphological profiling using multiplexed fluorescent dyes. Nat. Protoc. 11, 1757–1774 (2016).

